# Evidence for high intergenic sequence variation in heterozygous Italian ryegrass (*Lolium multiflorum* Lam.) genome revealed by a high-quality draft diploid genome assembly

**DOI:** 10.1101/2021.05.05.442707

**Authors:** Dario Copetti, Steven A. Yates, Maximilian M. Vogt, Giancarlo Russo, Christoph Grieder, Roland Kölliker, Bruno Studer

## Abstract

**Background:** Over the last decade, progress in DNA sequencing technologies and assembly methods allowed plant scientists to move beyond the use of model organisms and work directly on the genomes of the major crops. Forage grass research can also benefit from this revolution, enabling progress in population genetic studies, functional biology, and genomics-assisted breeding. Due to its large genome size and high repeat content, so far only incomplete and fragmented assemblies are available for the grasses of the *Lolium* and *Festuca* species complex.

**Findings:** Here, we report a highly contiguous draft assembly of Italian ryegrass (*L. multiflorum* Lam.), spanning 4.5 Gb and with a N50 of 3 Mb, containing ~70,000 gene models. Thanks to its relatedness to barley, 78% of the assembly was anchored on seven pseudomolecules. The high heterozygosity of the plant allowed obtaining a diploid assembly – i.e. across 95% of the assembly, both alleles were assembled on separate sequences. This feature allowed unraveling a very high amount of intergenic sequence variation between allelic sequences.

**Conclusions:** We present a nearly complete genome assembly of a genotype used in contemporary Swiss forage grass breeding programs. This work shows how genomic research has improved, allowing to decode the genetic code of large, complex, and heterozygous plants. It also allows the functional characterization of the ryegrass gene repertoire and the large-scale development of molecular markers. Furthermore, it paves the road for a reference-based characterization and exploitation of the genetic variation within the *Lolium* and *Festuca* species complex.

## Data description

### Context

Grasses of the genus *Lolium* are the most important forage and turf grasses worldwide [1]. Italian ryegrass (*Lolium multiflorum* Lam.), for example, provides an important feed source for roughage-based ruminant livestock production and is valued for its high palatability and biomass yield [2,3][3][3]. Italian ryegrass is cultivated in temperate regions, is naturally diploid (2n = 2x = 14), and characterized by a highly effective two-locus gametophytic self-incompatibility system that prevents self-pollination [4]. Such allogamous species maintain high levels of heterozygosity in their genomes and are often genetically improved as open-pollinated populations [5]. Both self-incompatibility and the breeding strategy have contributed to the high genetic variability that is present within as well as between ecotype, breeding populations, and commercial cultivars [6].

Like many grasses, members of the *Lolium* and *Festuca* species complex have large (above 2 Gb) genomes, varying in size and ploidy level [7,8]. The genome size of the economically most important species perennial ryegrass (*L. perenne* L.), Italian ryegrass and meadow fescue (*Festuca pratensis* Huds) are in a similar range and vary between 2.6 and 3.2 Gb [7]. Previous efforts at sequencing ryegrass genomes resulted in partial and fragmented assemblies [9–12], mainly due to the high amount of repeated sequences and the limitations of *de novo* assembly algorithms adopted. In addition, the two publicly available perennial ryegrass assemblies originated from inbred and highly homozygous genotypes. On the one hand, such highly inbred genotypes facilitated the assembly process, but, on the other hand, they are often not representative of the advanced germplasm commonly used in breeding programs. Though highly contiguous, the chromosome-scale genome assembly of orchardgrass (*Dactylis glomerata* L.) [13] is devoid of any allelic sequence and the species belongs to a different subtribe (Dactylidinae) than *Lolium* and *Festuca* (Loliinae).

Despite *de novo* genome assembly methods are being developed since more than two decades [14], only recently the community started focusing on highly heterozygous genomes [15–21]. Typically, high amounts of sequence variation between alleles prevent contig extension and scaffolding. Therefore, assemblies of heterozygous organisms result in fragmented (for the frequent branching of the assembly graph) and redundant (with divergent allelic regions assembled separately) reconstructions [22]. The ideal assembly of a heterozygous species should instead present both alleles separately (referred to as being diploid), span the whole genome (being complete), and do it in the smallest possible number of sequences (being contiguous).

Diploid-aware genome assembly algorithms were first developed for short read data [22–24]. However, due to limited read length and genome complexity, only a (often small) fraction of the alternative allelic sequence could be reconstructed. The 10x Genomics’ Chromium method offers another solution for generating diploid assemblies. By labeling separately large DNA fragments, this linked read technology has the potential to reconstruct entire diploid genomes in an allele-specific manner [25]. However, convincing examples of diploid and truly phased assemblies produced with such platform are still lacking. A proprietary method for *de novo* genome assembly from short read data was developed by NRGene (Ness-Ziona, Israel) and, among others, several assemblies of plant genomes are available [26–30,18]. In the case of the heterozugous apple genome, the algorithm successfully assembled the two homologous regions in to separate sets of sequences [20]. Here, we present the diploid assembly of a diploid Italian ryegrass genome. The high degree of heterozygosity allowed us to reconstruct in two allelic sequences the vast majority of the genome, resulting in the first nearly complete diploid assembly of a forage grass species.

## Materials, methods, and results

### Plant material

The plant material used in this study was obtained by germinating a single seed of the Italian ryegrass cultivar Rabiosa (the genotype was called M.02402/16) and multiplying it clonally through tillers. The cultivar Rabiosa was developed within the Fodder Plant Breeding Program of Agroscope, Switzerland and represents contemporary elite germplasm. It entered the Swiss national list of recommended varieties in 2015 as best variety, showing high yields, best persistence and very good resistance against bacterial wilt caused by *Xanthomonas translucens* pv. *graminis* [Additional file 1].

### Cytogenetics

For this and the following sections, a more detailed description of the methods, tools, and parameters is available in an additional document [Additional file 2]. Prior to sequencing, the ploidy level of M.02402/16 was evaluated by counting the chromosomes. Root tips were soaked in a 3:1 solution of ethanol and acetic acid at 4 degrees. After washing, the tips were soaked in 1 N HCl at 60 °C for 10 min before staining with Schiff reagent for 1 h. The tips were then transferred to glass microscope slide and squashed.

The direct observation of sets of fourteen chromosomes in multiple nuclei confirmed the diploid nature of M.02402/16. Interestingly, four supernumerary elements with no visible or acentric centromeres were consistently observed (Suppl. Figure 1 a). Possibly, these fragments are the result of chromosomal fragile sites, which are 45S rDNA carrying chromatin fibers, connecting the fragments with centromeric chromosomes [31].

### Nucleic acid extraction and sequencing

To obtain an accurate reconstruction of the genome, we sequenced at high depth three different types of short-read libraries (Suppl. Table 1). DNA was extracted from young leaves of M.02402/16 following a modified CTAB protocol. Sequencing libraries were produced following NRGene’s standard *de novo* whole-genome shotgun (WGS) and 10x Genomics Chromium protocols. Sequencing mode and depth varied according to the library type. The overall coverage was above 500 genome equivalents of a ~2.5 Gb haploid genome. The mRNA sequence data used for genome annotation was derived from the clonal propagations of M.02402/16. Four tissue types were sampled: mature leaves, apical meristems, roots, and flowers. RNA-Seq libraries were sequenced with the Illumina platform with a read length of 2x150 bp. All the raw data generated was deposited in GenBank SRA archive under the accession number SRP151232 (Suppl. Table 2).

### Nuclear genome size estimation and assembly

To estimate *in silico* the genome size from the sequencing data, estimate_genome_size.pl [32] was run with the raw paired-end reads and a k-mer size of 17. The haploid genome was estimated to be 2.464 Gb large, comparable to the 2.567 and 2.661 Gb estimated by flow cytometry on closely-related species [7,33]. The k-mer distribution plot revealed the presence of two peaks (Suppl. Figure 1 b). The peak around k-mer frequency 40 was composed of heterozygous k-mers, while the second peak (mode at ~80) had the homozygous k-mers, i.e. sequences identical between the two alleles. Though some heterozygosity was expected due to the obligate outcrossing nature of the species, the dominance of sequences in a heterozygous state (about two-thirds of true low-copy k-mers) was unanticipated. The ~1.34 Tb of raw sequencing data were assembled using NRGene’s DeNovoMAGIC^™^ 3.0 pipeline, a proprietary algorithm that has shown to produce highly-contiguous assemblies of large and repeat-rich plant genomes [18,27–30]. The initial assembly consisted of 227,513 scaffolds, spanning more than 4.53 Gb, and containing 1.43% Ns. Half of the assembly was contained in the 421 longest scaffolds (L50), whose length was greater than 3.05 Mb (N50, Table 1). The GC content was 43.2%, a typical value for a grass genome [34]. By size and contiguity metrics, this assembly is orders of magnitude greater than previously published *Lolium* assemblies [9,12] and comparable to others obtained with the same method (Suppl. Table 3 [27,30,35]).

**Table 1.**
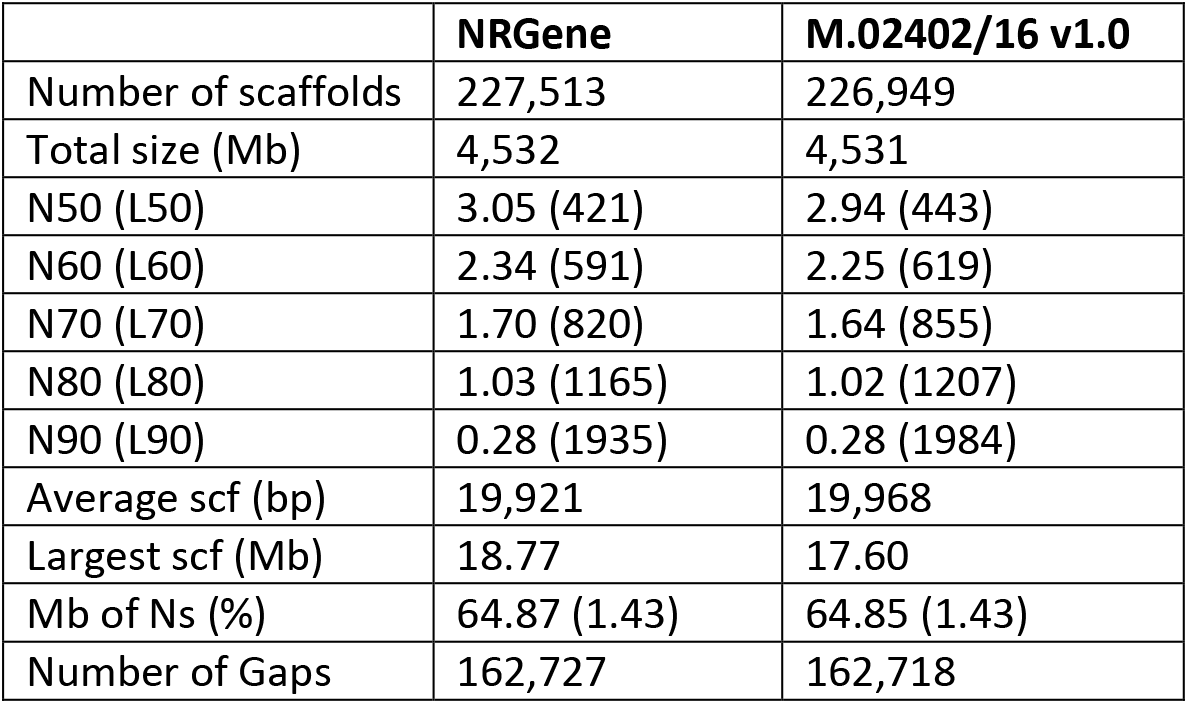
Statistics of the M.02402/16 assembly upon delivery from NRGene and at its final curation stage.

### Assembly quality control and sequence anchoring

Scaffolds were inspected for the presence of chimeric sequences by leveraging on the collinearity with barley (*Hordeum vulgare* L.) and on the availability of a genetic linkage map of the closely related species *L. perenne*. Details of the method are reported in [Additional file 2]. In short, the following four sets of evidence were deployed: i) *Lolium* expressed sequence tags (ESTs) and Unigenes were aligned with GMAP (v. 2017-09-30 [36]) to the M.02402/16 assembly and to the barley proteome. Eventually, 4,579 alignments linked 1,070 scaffolds to barley genes ([Additional File 3], sheet Unigenes). ii) Provisional Rabiosa gene models were derived from *L. perenne* RNA-Seq data (Studer *et al*., unpublished) with MAKER (v. 2.31.9 [37]) and aligned as above against barley genes. After parsing, 7,082 gene alignments connected 1,435 M.02402/16 scaffolds to anchored barley gene models ([Additional File 3], sheet RNA-Seq). iii) To identify collinear blocks of three or more genes, the M.02402/16 assembly was aligned to the barley assembly, identifying 631 collinear barley-ryegrass scaffolds ([Additional File 3], sheet SynMap). Lastly, iv) the 762 markers of the perennial ryegrass genetic linkage map [38] were aligned with BLASTN (v. 2.5.0 [39]) to the assembly. Using this genetic evidence from a closely related species, the genetic markers anchored 411 scaffolds to *L. perenne* linkage groups (LGs, [Additional File 3], sheet Pfeifer 2013). Overall, the presence of many blocks of collinear genes between barley and Italian ryegrass scaffolds confirms the high conservation of the gene order between the two species (Figure 1 a). Furthermore, the mapping of pairs of M.02402/16 paralogs to a single barley gene supports the structure of the scaffolds and the diploid nature of the assembly (Figure 1 b).

**Figure 1.**
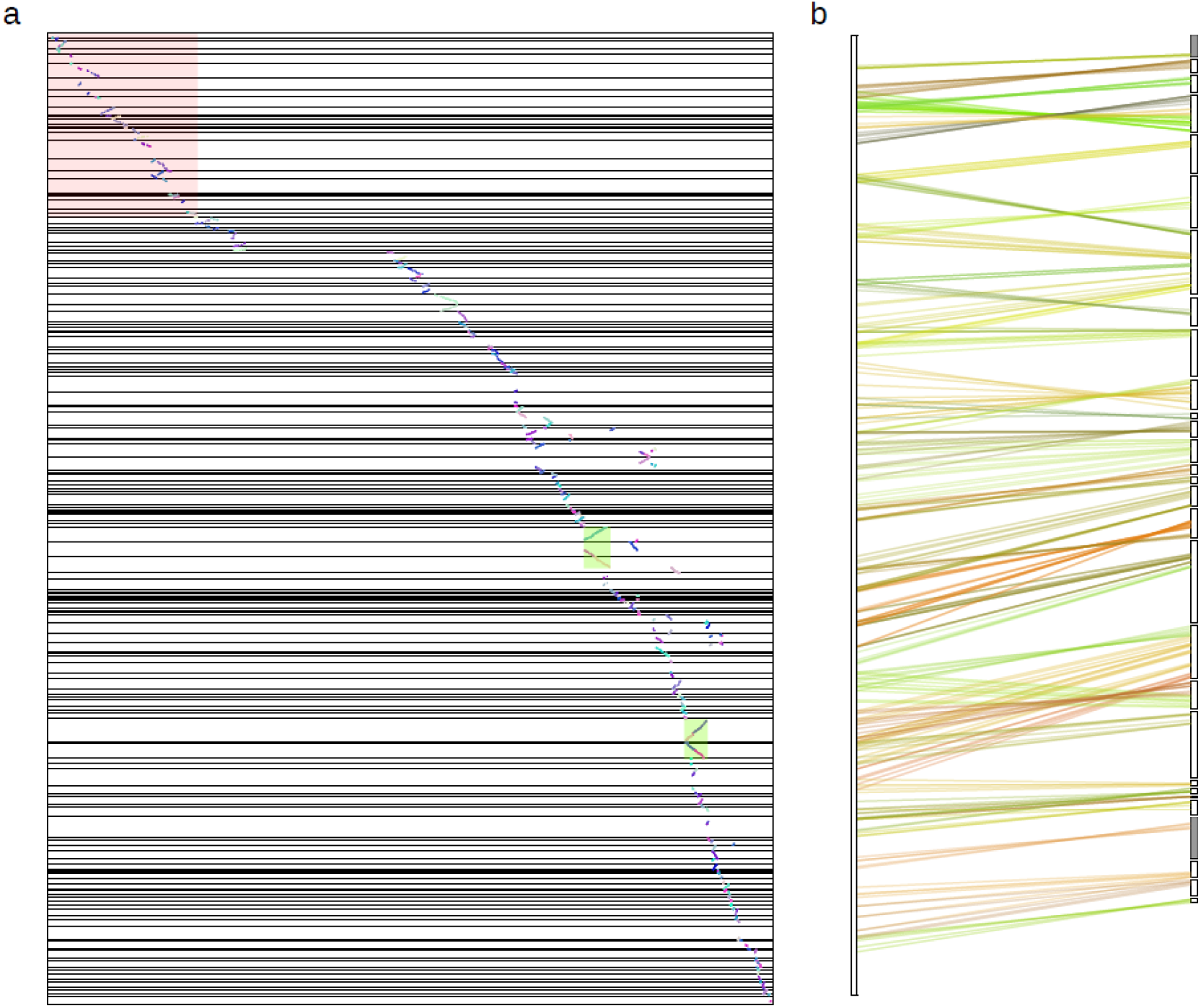
Collinearity of barley-M.02402/16 scaffolds and allelic redundancy. (a) Alignment of M.02402/16 scaffolds (y axis) to barley chromosome 1 (x axis). Collinear blocks are depicted as chains of colored dots, green rectangles highlight examples of collinear blocks of allelic scaffolds. The coordinates of both axes are expressed in gene counts. The area within the red rectangle is magnified in (b), displaying M.02402/16 scaffolds (bars on the right) that align with the first 120 Mb of barley chromosome 1 (left bar). Multiple pairs of allelic ryegrass scaffolds show hits (differently colored lines denote different scaffolds) to the same barley genes. The two gray scaffolds are the only cases where, for the barley homologue, the second collinear scaffold could not be found.

These four analyses produced a set of scaffolds that had stretches of three or more genes aligning to two different barley chromosomes. With additional evidence of steep dips in coverage of both MP and 10x Chromium data, 43 of these were split using bedtools v. 2.26.0 [40]) in two or three parts. This edited version of the assembly was called Rabiosa M.02402/16 v1.0.

The assembly maintained high contiguity (half of it is in scaffolds of 2.94 Mb or longer, and less than 2,000 scaffolds representing the longest 90% of the assembly, Table 1), an important feature that improves significantly the quality of the available genomic resources of the *Lolium* and *Festuca* species complex [9,10,41]. In comparison with other publicly available genome sequence resources of ryegrasses, the M.02402/16 assembly shows a >40-fold improvement in contiguity (Suppl. Table 3) and a N50 value similar to other well assembled (yet large) grass genomes [27,42,43]. Similarly, all the public ryegrass assemblies span just a minority of the estimated genome size (Suppl. Table 3). Of notice, the N50 of this assembly is more than 4-fold larger than maize’s PH207 assembly – an inbred line of comparable genome size (2.6 Gb), obtained with the same method [29]. This discrepancy supports the evidence that the genome composition (e.g. amount and type of TEs that proliferated) directly affects the quality of an assembly.

### Completeness

To assess the completeness of the M.02402/16 assembly with respect to plant ultra-conserved single-copy orthologs (SCOs), a BUSCO (v. 3 [44]) analysis was performed. The run identified 96.8% of the expected near-universal single-copy orthologs, with only 1.3% being fragmented and 1.9% missing. Though the analysis considers only ultraconserved single-copy genes, the completeness of the M.02402/16 assembly was higher compared to other ryegrass assemblies and comparable to other key grass reference sequences (Suppl. Figure 2). Importantly, 69.7% of the complete genes were present in two copies, only 21.9% in single copy and 5.1% had three or more copies. Among plant genomes, only the bread wheat [30] and Wild Emmer wheat [27] assemblies show a similarly high fraction of SCOs present in multiple copies. This latter evidence is, however, consequence of the ploidy level of the species, rather than the presence of both alleles in the same assembly, as is the case for M.02040/16 (Suppl. Figure 2).

To estimate the fraction of protein-coding genes present in the assembly, raw RNA-Seq reads were aligned. The read mapping rate ranged between 70% and 97%, with lower rates for the perennial ryegrass samples (Suppl. Table 2). M.02402/16 flower, leaf, and meristem transcripts aligned at higher proportion, between 95.6% and 97.4%, while the root samples had a slightly lower mapping rate of around 90%. Upon *de novo* assembly of the same data with Trinity (v. 2.5.1 [45]), about 97% of flower, leaf, and meristem unigenes aligned to the reference, while less than 58% of the root transcripts had a hit. Alignment of a subset of these unmapped transcripts to NCBI non-redundant database showed that these root transcripts belong mostly to fungal and bacterial proteins, thus may represent contamination of the sample with soil-borne organisms. Unmapped hits from the other three organs had mostly partial hits to predicted grass proteins. Overall, the high alignment of transcripts indicates a near complete representation of the transcribed portion of the genome in the assembly.

Aligning WGS reads to an assembly can help determining whether the reference (or a region of it) was reconstructed from one or both alleles of the genome, with different coverage depths pinpointing to sequences representing one or both alleles. To identify the extent to which the assembly collapsed (i.e. is haploid), all paired-end (PE) reads were aligned to the final M.02402/16 scaffolds using BWA-MEM (v. 0.7.17 [46]). Overall, 99.8% of the reads aligned to the reference and 89.4% of these aligned as proper pairs. Of the reference bases, 99.7% were covered by at least one read, and only 1.1% of the assembly had a coverage lower than 10 genome equivalents. The high alignment rate and breadth of coverage show a very complete representation of the sequenced space in the assembly.

In a genome assembled with a consistent ploidy level, the read coverage should show a unimodal distribution and be in agreement with the assembly ploidy and sequencing depth. The alignment of the above PE data to the assembly presented a single peak with mode at 57× coverage, and the two interquartiles at 37× and 66× (Figure 2 a). The more prominent shoulder at lower coverage (<30×) was consequence of the low mappability of PE data in repetitive regions. At higher coverage values, a minor shoulder was present approximately at a value twice the mode. This small fraction of reference with higher coverage (arbitrarily confined between 90<×<150) represented assembly regions where the two haplotypes were collapsed, and it contained about 7.4% of the total area between coverage 1× and 200×.

**Figure 2:**
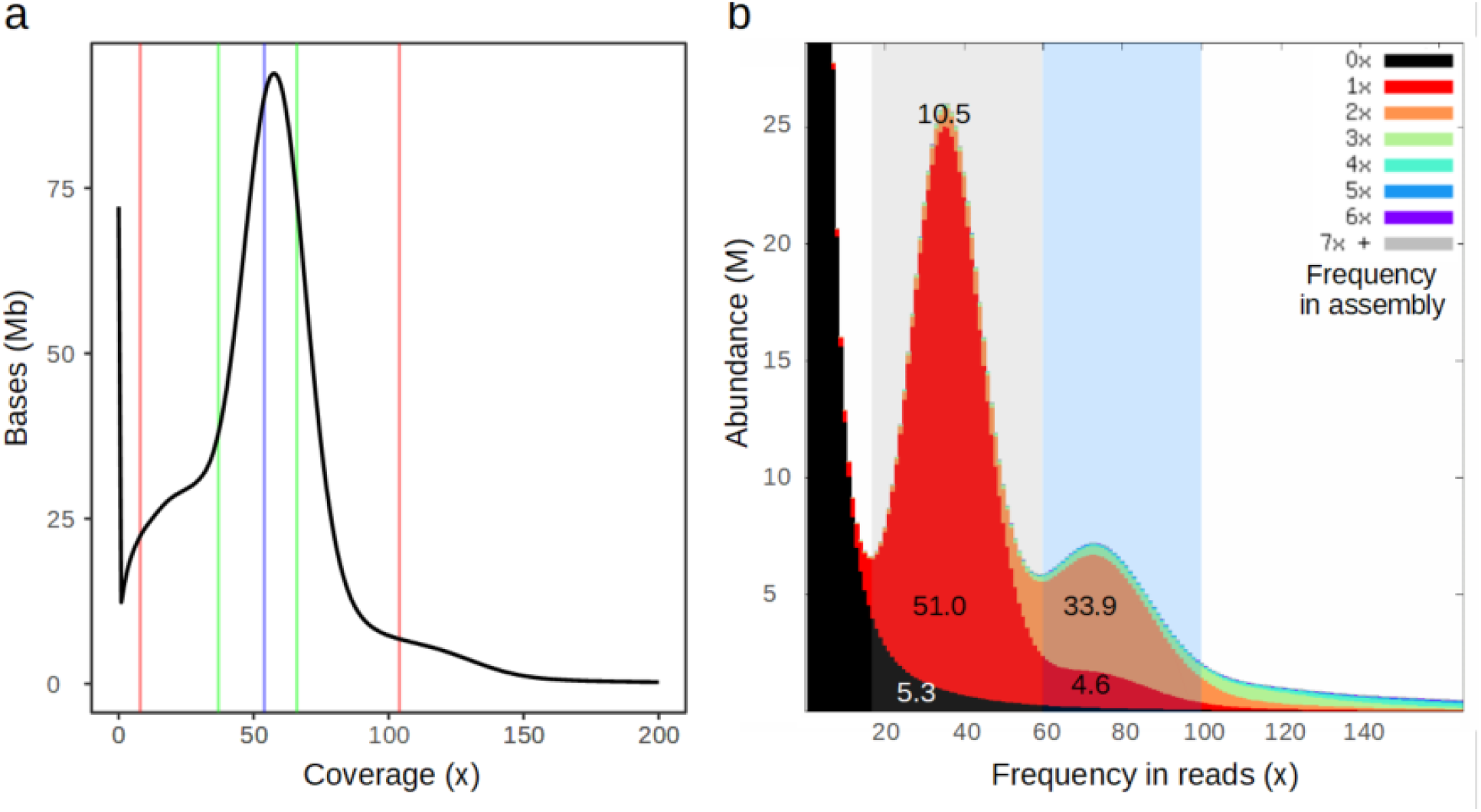
Estimation of the haploid and diploid fractions of the assembly. (a) Coverage distribution of about 115× PE reads to the M.02402/16 assembly. The presence of a main peak around half of the expected coverage for a haploid genome implies that most of the assembly is diploid – i.e. that most of the regions in the genome are assembled in two separate sequences. The distribution was calculated for all bases after removing the few counts with more than 200× coverage. The blue bar indicates the median (at 54×), the green lines are the 25^th^ and 75^th^ quartiles (37× and 66×, respectively), the red lines are the 5^th^ and 95^th^ quartiles (8× and 104×). (b) KAT plot comparing presence and abundance of 23-mers between reads and assembly. On the x axis, the k-mer frequency in the raw reads, on the y axis, abundance of the k-mers in the assembly (split by their abundance in the different colors). The k-mer profile of the reads was split in two areas, representing the interval of the heterozygous and homozygous peak (gray and blue shaded areas, respectively). The numbers indicate the fraction of the sequence that was missing from the assembly (5.3%) or present at different abundances (other values).

The K-mer Analysis Toolkit (KAT) was used to assess the completeness of the M.02402/16 assembly with respect to the information present in the raw reads (Figure 2 b). The analysis highlighted that: i) almost two-thirds of the k-mers in the assembly are heterozygous (the proportion of the gray and blue shaded intervals is 61.5% and 38.5%, respectively). ii) If they could be assembled, the unincorporated k-mers would increase the assembly of only 5.3% (sum of the black area between ×=17 and 100). iii) The amount of assembly artifacts is very low. About 10% of the heterozygous k-mers are present two or more times in the assembly (while they should be present only once) and 0.003% of the k-mers of the assembly are not found in the raw reads (novel sequences generated during the assembly). iv) Regions homozygous in the genome but collapsed into one copy in the assembly span only 4.6% of the k-mers (red area under the blue shading). This fraction represents only about 12% of the homozygous k-mers, implying that most of the identical allelic regions are assembled separately in the assembly. The highly similar proportion of sequenced and assembled heterozygous k-mers (67.1% and 61.5%, Suppl. Figure 1 and Figure 2 b, respectively) indicates a very good representation of the diploid sequenced space in the assembly. The two different k-mer analyses support the evidence that the M.02402/16 genome is highly heterozygous, and that the assembly contains the vast majority of the sequence variation, correctly allocated to allelic sequences.

### Anchoring scaffolds to chromosome pseudomolecules

With the exception of a translocation involving chromosomes 4 and 5 [38,47], ryegrass and barley show high synteny. Taking advantage of the collinearity with barley and the alignment of the *L. perenne* genetic map, M.02402/16 scaffolds were assigned, ordered and oriented into *L. multiflorum* pseudochromosomes. To account for the translocation of the terminal part between barley chromosome 5 and ryegrass chromosome 4, the scaffolds aligning to gene models HORVU5Hr1G092630-HORVU5Hr1G116580 (corresponding to *L. perenne* markers PTA.275.C1 and PTA.456.C3, respectively [38]) were moved to the top of linkage group 4 (LG4) in reverse order. The four lines of evidence were used as input in ALLMAPS [48] and scaffolds with ambiguous alignments or aligning to less than two genes belonging to the same region were left unanchored. The output in agp format is at [Additional file 3], sheet ALLMAPS. The result highlights a good correlation between barley gene order (or *L. perenne* genetic order) and the placement of the M.02402/16 scaffolds and their markers therein, both at the whole-chromosome level and between markers within scaffolds (Figure 3).

**Figure 3:**
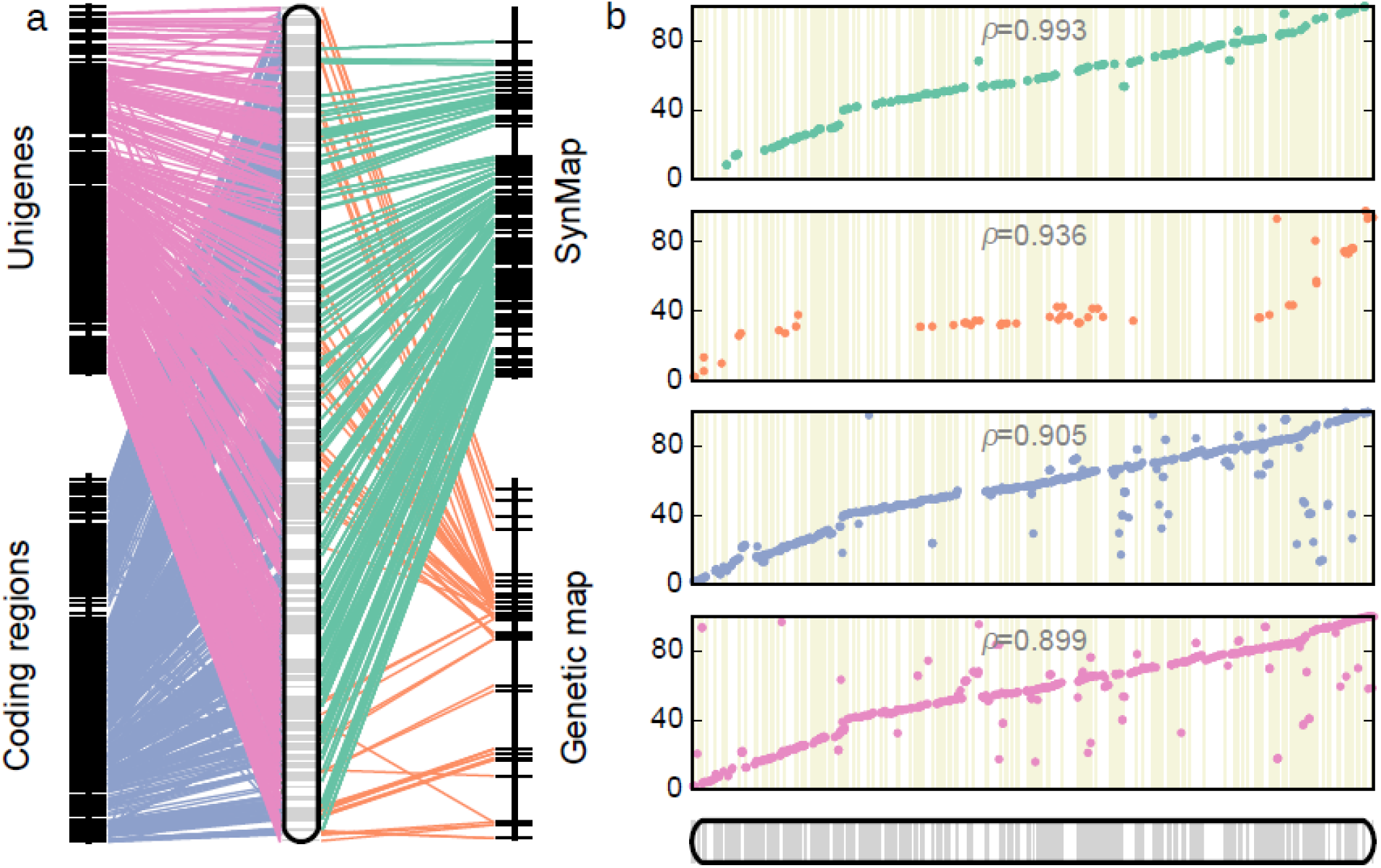
Assignment of the M.02402/16 assembly to chromosome pseudomolecules. (a) Contribution of each set of evidence to assign, order, and orient scaffolds on pseudochromosome 1. Lines connect a marker (genic or genetic) from its relative position on barley to the corresponding ryegrass scaffold. (b) Correlation between physical position on the ryegrass pseudomolecule (x axis) and the relative position of the barley gene order (y axis). The four sets of evidence show high consistency as well as complementarity.

Overall, 3.5 Gb of the assembly (77.7%, represented in 1,734 scaffolds) were assigned to a M.02402/16 pseudochromosome, with the majority (70% of the scaffolds) also being oriented (Table 2). The N50 of the anchored scaffolds was 3.75 Mb, while the ~1 Gb that was left unplaced was mostly composed of shorter sequences (N50 0.47 Mb). These sequences did not show clear collinearity with the barley assembly, nor contained genetic markers.

**Table 2:**
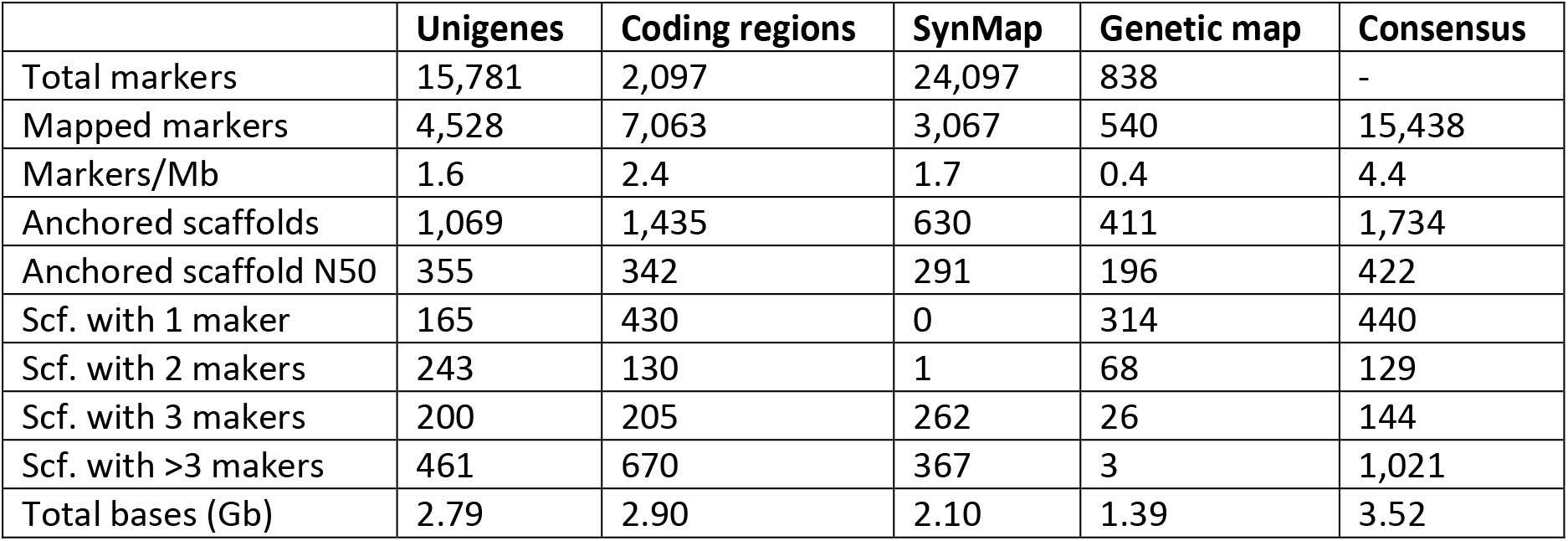
Results of the assignment of M.02402/16 scaffolds to a *L. multiflorum* chromosome pseudomolecule.

|  | Unigenes | Coding regions | SynMap | Genetic map | Consensus |
| --- | --- | --- | --- | --- | --- |
| Total markers | 15,781 | 2,097 | 24,097 | 838 | - |
| Mapped markers | 4,528 | 7,063 | 3,067 | 540 | 15,438 |
| Markers/Mb | 1.6 | 2.4 | 1.7 | 0.4 | 4.4 |
| Anchored scaffolds | 1,069 | 1,435 | 630 | 411 | 1,734 |
| Anchored scaffold N50 | 355 | 342 | 291 | 196 | 422 |
| Scf. with 1 maker | 165 | 430 | 0 | 314 | 440 |
| Scf. with 2 makers | 243 | 130 | 1 | 68 | 129 |
| Scf. with 3 makers | 200 | 205 | 262 | 26 | 144 |
| Scf. with >3 makers | 461 | 670 | 367 | 3 | 1,021 |
| Total bases (Gb) | 2.79 | 2.90 | 2.10 | 1.39 | 3.52 |

The importance of each dataset in assigning scaffolds to pseudomolecules is emphasized by the fact that, taken individually, the four sets of evidence anchored more than twice as many scaffolds – implying that there is limited redundancy between the inputs.

### Repeat annotation

Due to the lack of exemplar TEs and repeats isolated from forage grass genomes, RepeatExplorer (v. v0.2.8-2479 [49]) as in [50] and RepeatModeler [51] were used to develop two repeat libraries. As expected, TEs and repeated sequences cover more than three quarters of the assembly, and like in most plants, LTR-RTs are the dominant fraction (two thirds of all TEs/repeats and half of the assembly size, Suppl. Table 4). The high amount of highly-repetitive sequence was also evident from a sequence dotplot [52] of two allelic scaffolds, where LTR-RT or satellite repeats (Suppl. Figure 3, within red and blue rectangles, respectively) occupy most of the region. Furthermore, only small and scattered tracts are shared between alleles (short diagonal lines highlighted by green triangles in Suppl. Figure 3), and the intervening sequence is highly polymorphic (deviation of the lines from the expected continuous diagonal).

### Protein-coding genes and non-coding RNA annotation

By combining different types of evidence, the MAKER (v. 2.31.9 [37]) pipeline identified 70,688 protein-coding loci and 99,288 splice variants. The gene space covered only 6.6% of the assembly and the exons and introns had a mean length of 310 bp and 628 bp, respectively. The average gene spanned 4.1 kb and the average protein product was 347 aa. Overall, the consistency of the final gene models to the input evidence was very high, with about 52% of the isoforms having an annotation edit distance (AED) score below 0.2 and more than 95% below 0.5. The vast majority (90.2%) of the predicted protein-coding loci were located on the 1,743 scaffolds (spanning almost 80% of the assembly bases) that were assigned to a pseudochromosome.

The protein-coding genes were functionally annotated by alignment (BLASTP) to Arabidopsis genes (chosen for the higher number of experimental annotation and its manual curation) and by retrieving the functional annotation with InterProScan (v.5.31-70.0, using default setting and databases [53]). Finally, a non-redundant list GO terms was created from both the Arabidopsis and InterProScan sources. Of the 99,288 proteins searched, 66,533 (67%) were assigned an annotation, of which 59,469 (60% of the total) were assigned at least one GO term.

By surveying the proportion of expected ultraconserved single-copy orthologs and conserved gene families, the M.02402/16 annotation had comparable completeness as other highly-curated assemblies (e.g. barley, *Brachypodium*, rice [35,42,54]), and showed a clear improvement compared to the available *L. perenne* assembly (Suppl. Table 5). While BUSCO (v. 3 [44]) scans the annotation for the presence of evolutionary conserved single-copy orthologs, the coreGF (v. 2.5 [55]) completeness score reflects the presence of highly-conserved gene families in a specific plant clade – in this case 7,076 genes conserved across all monocots. Therefore, though the perennial ryegrass’ BUSCO score is quite high (87% of complete SCOs vs 94% in M.20402/16), the assembly is missing a good amount of multi-copy genes. The 75% coreGF score is a rather low value compared to the other grass assemblies. Importantly, the M. 02402/16 gene annotation retains the high amount of duplicated SCOs that was already detected at the assembly level, providing support that the high number of genes (twice as many as a typical diploid grass species) is indeed allelic redundancy (Suppl. Table 5). Considering the presence of both alleles for most regions of our assembly, the >70,000 genes predicted are a value in line with previous observations in other diploid grasses, ranging from ~32,000 in bamboo [56], to 35,000-40,000 in rice [54,57] to 39,000 in barley [35], to ~40,000 in maize and *A. tauschii* [28,43]. Compared to the about 28,000 gene models predicted in *L. perenne* [9], the current assembly significantly broadens the gene catalog from forage grass genome data.

### Sequence diversity in allelic regions

To draw a more qualitative representation of allelic regions in the M.02402/16 genome, we investigated the region surrounding the *Ethylene Overproducer 1-like (ETO1)* gene, which regulates ethylene production by negative inhibition of the enzyme 1-aminocyclopropane-1-carboxylate synthase (ACS) – the rate limiting step in ethylene production [58]. The *L. multiflorum* orthologs (corresponding to the rice sequence 015628339.1, in LOC_Os03g18360 on *O. sativa* vg. *Japonica* [54]) were identified by GMAP (v. 2018-03-25, [36]) with gene models Lmu01_16T0000470 and Lmu01_1530T0000400, located on scaffold 16 and scaffold 1530, respectively. The two scaffolds were anchored on chromosome 4, and showed high collinearity with the region at 460-474.5 Mb on barley chromosome 4H (delimited by genes HORVU4Hr1G055050 and HORVU4Hr1G056500, respectively, Figure 4 a [35]). Thirty-five barley genes in that region have hits to one or both M.02402/16 scaffolds. The common region surrounding *ETO1* on the two allelic M.02402/16 scaffolds spans approximately 3 Mb on each allele. A dotplot alignment of this region in the two M.02402/16 scaffolds shows that, between two alleles, the majority of the shared sequence contains genes, with most of the intergenic space being allele-specific (interruptions and shifts of the diagonal, Figure 4 b). In terms of sequence similarity, the alignment (MUMmer4.x [59]) of the allelic region revealed that less than 40% of the nucleotidic sequence could be aligned in a 1-to-1 relationship (Suppl. Table 6), with 3.2% mismatches (or allelic single-nucleotide polymorphysms, SNPs) and 1.3% insertions/deletions (indels). On the *L. perenne* P226/135/16 assembly [9], the *ETO1* gene was assembled in a ~70 kb scaffold, and the same barley region surrounding it contained at least 32 P226/135/16 scaffolds. A dotplot analysis of the region in the two assemblies still showed only interspersed microcollinearity, implying high intergenic sequence variation between the two *Lolium* species (Suppl. Figure 4). Quantitatively, only 17.1% of the *L. perenne* sequence could be aligned to the *L. multiflorum* assembly (covering only 13.9% of the region in M.02402/16), with 3.2% SNPs and 2.0% indels (Suppl. Table 6).

**Figure 4:**
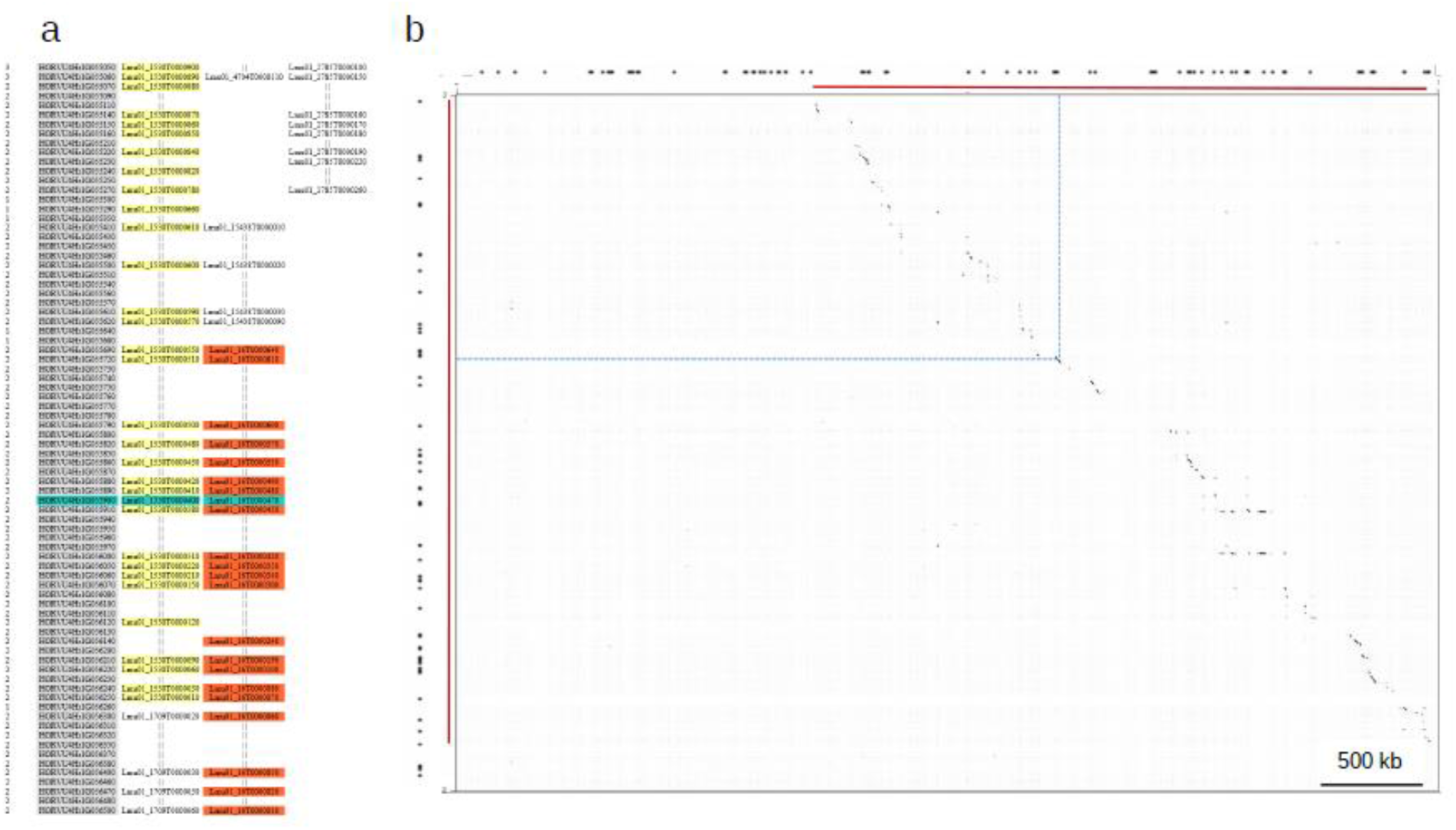
Collinearity of the diploid assembly with barley around the *ETO1* locus and allelic variation. (a) Alignment of the M.02402/16 assembly around the barley *ETO1* locus. Each row represents the position of a barley gene, and orthologous genes between barley (gray shaded column) and Italian ryegrass genes (three rightmost columns) lie on the same row. Genes of scaffold 1530 and scaffold 16 collinear with barley genes surrounding *ETO1* (position highlighted in green) are highlighted in yellow and orange, respectively. Genes not highlighted are from flanking scaffolds that do not contain the *ETO1* ortholog. The numbers on the left report the syntenic depth at each reference gene. (b) The dotplot alignment of scaffold 1530 (x axis) and scaffold 16 (y axis) shows the low amount of shared sequence between allelic scaffolds. Coordinates increase from the top left corner. Outside the plot, red bars span the region of overlap between the two scaffolds; the black dots are the predicted genes, the dashed blue line projects the coordinates of *ETO1* on the plot. For clarity, one of the two sequences was hard-masked to remove the high background signal that interspersed repeats generate when present in multiple copies on both sequences.

The amount of intergenic sequence variation observed in M.02402/16 at the same multi-Mb scale was further compared to other grass genomes. About two-thirds of the sequence of two maize lines (B73 and PH207, belonging to different heterotic groups, [26,60]) was shared, with less than 1% SNPs and indels (Suppl. Table 6 and Suppl. Figure 5 a). Comparison between rice species [54,61] provided metrics more close to what was observed between ryegrass sequences: only about 17% of the sequence was shared (with at least 4.5% SNPs and 1.9% indels) between cultivated Asian rice and *Oryza brachyantha*. The two rice species diverged ~15 million years ago [62] (Suppl. Figure 5 b). Though purely descriptive, the dotplot comparison between different individuals or species reveals an unprecedented amount of intergenic sequence variation between ryegrass alleles, depicted by the loss of sequence homology between allelic sequences.

### Summary and conclusions

Aiming to establishing a high-quality reference genome assembly for forage and turf grass research, we sequenced and assembled a heterozygous individual (M.02402/16) of the Italian ryegrass cultivar Rabiosa. Extensive and complementary validation analyses of the sequenced space and gene content supported the high quality and diploid nature of the M.02402/16 assembly. Both the assembly size and the number of genes are close to values expected for a diploid genome. Also, the diploid status is confirmed by the read coverage distribution and by pairs of ryegrass scaffolds that are collinear with barley regions. The unanticipated feature that this genome unraveled was the high amount of intergenic large-scale sequence variation (not due to SNPs) observed between allelic sequences. Immediate applications of this resource include: the establishment of modern functional and comparative genomic analyses in the realm of forage grasses, its use as a reference to anchor assemblies of closely-related varieties, as a reference for resequencing projects, marker discovery, and the availability of an exhaustive catalogue of ryegrass genes.

Even though highly dynamic at the genomic level, the ryegrass species are still interfertile and can produce viable offspring, even with the closely-related *Festuca* spp. [63–65]. This allows to combine properties in allopolyploids or transfer with ease traits between species, as demonstrated in current forage and turfgrass breeding programs [66,67]. To raise such a powerful system to a modern genomics standard, the development of a pan-genome project encompassing multiple forage and turfgrass species will be a key step (Studer *et al*., in progress). With the present draft, we demonstrate the possibility to obtain a mostly complete and diploid draft assemblies for outbreeding plant species with a large and complex genome.

## Availability of Data and Materials

The data produced for this project has been deposited to NCBI database under the BioProject NNNNNNN. The raw data and the assembly are available under the accession NNNNNNNN and NNNNNNNNN, respectively. The genome annotation is available at DOI:NNNNNNNNNN.

## Additional files

**Additional File 1. Description of the development of the Rabiosa variety.**

Brief description of how the Rabiosa variety was obtained.

**Additional File 2. Detailed description of material and methods.**

For brevity, the main manuscript contained the most salient methods necessary to support our findings. The complete description of methods and software is reported with full detail in this file.

**Additional File 3. Anchoring of M.02402/16 scaffolds.**

Each sheet lists the parsed alignments that were kept for identifying chimeric scaffolds and anchoring/orienting scaffolds in pseudomolecules. The ALLMAPS sheet contains the .agp file, output of ALLMAPS.

**Supplementary_Tables.xls**

**Supplementary_Figures.pdf**

## List of abbreviations

AED: annotation edit distance
CDS: coding sequence
EST: expressed sequence tag
GEM: gel beads in emulsion
indel: insertion/deletion
LG: linkage group
LTR-RTs: long terminal repeat-retrotransposon
MP: mate pair
PE: paired end
SCOs: single copy orthologs
SNP: sinlge-nucleotide polymorphism
TE: transposable element
WGS: whole-genome shotgun

## Consent for publication

Not applicable

## Author’s contribution

S.Y. and B.S. designed the sequencing project, C. G. developed the cultivar Rabiosa, S.Y. and R. K. extracted nucleic acids, D.C., M.M.V., G.R., and S.Y. performed the analyses, D.C., S.Y., R.K., and B.S. wrote the manuscript. All authors approved the final version of the manuscript.

## Acknowledgements

The authors would like to thank Beat Boller for the initial germplasm collection, Roberto Copetti and Jon Wright for their insights on the methods, NRGene for scientific support, and the Arizona Genomics Institute for making available computing resources.

---

**Suppl. Table 1. Statistics of the raw DNA sequencing data produced for the genome assembly.**

**Suppl. Table 2. Features of the RNA-Seq data produced and used in this study.**

**Suppl. Table 3. Comparison of the statistics of publicly available assemblies for the Poaceae family**. Highly contiguous and complete assemblies are available only for agriculturally relevant species (wheat, barley, maize) or model species (*Brachypodium*), while all public ryegrass assemblies are either incomplete or fragmented. The M.02402/16 assembly introduces a reference-grade resource for the forage and turfgrass community.

**Suppl. Table 4: Diversity and abundance of repeated sequences and transposable elements in the M.02402/16 assembly.**

**Suppl. Table 5. Comparison of the completeness of the gene annotation in published grass genomes.** The coreGF score represents the % of coreGF families found in the annotation (7076 genes in the PLAZA v2.5 monocot dataset). BUSCO was run in protein mode, with Plantae as the SCO lineage and wheat as the species for Augustus gene models (only rice and sorghum were run with rice and maize species, respectively). The values for M.02402/16’s transcriptome are derived from the *L. multiflorum* RNA-Seq gene models that were one of the evidence for the MAKER annotation.

**Suppl. Table 6: Sequence alignment of ~3 Mb homologous regions in grasses.** The alignment between the two M.02402/16 scaffolds at the ETO1 locus was made only with the shared portion (red lines in figure ETO1 B). The *L. multiflorum/L. perenne* alignment was made between the whole scaffold 16 and the 32 P226/135/16 scaffolds that aligned in the region. For the alignments between *Zea mays* inbreed lines and rice species, the first 3 Mb of chromosome 1 of each assembly were aligned.

## References

1. Gilliland TJ, Johnston J, Connolly C. A review of forage grass and clover seed use in Northern Ireland, UK between 1980 and 2004. Grass Forage Sci. 2007; doi: 10.1111/j.1365-2494.2007.00588.x.

2. Bernard JK, West JW, Trammell DS. Effect of replacing corn silage with annual ryegrass silage on nutrient digestibility, intake, and milk yield for lactating dairy cows. J Dairy Sci. 2002; doi: 10.3168/jds.S0022-0302(02)74307-5.

3. Andrzejewska J, Contreras-Govea FE, Berzaghi P, Albrecht KA. Forage Accumulation and Nutritive Value of Italian Ryegrass–Kura Clover Mixture in Central Europe. Crop Sci. 2018; doi: 10.2135/cropsci2017.05.0274.

4. Cornish MA, Hayward MD, Lawrence MJ. Self-incompatibility in ryegrass. Heredity. 1979; doi: 10.1038/hdy.1979.63.

5. . Fodder Crops and Amenity Grasses. Ryegrasses. New York: Springer-Verlag; p. 211–60.

6. Kölliker R, Stadelmann FJ, Reidy B, Nösberger J. Genetic variability of forage grass cultivars: A comparison of Festuca pratensis Huds., Lolium perenne L., and Dactylis glomerata L. Euphytica. 1999; doi: 10.1023/A:1003598705582.

7. Kopecký D, Havránková M, Loureiro J, Castro S, Lukaszewski AJ, Bartos J, et al.. Physical distribution of homoeologous recombination in individual chromosomes of Festuca pratensis in Lolium multiflorum. Cytogenet Genome Res. 2010; doi: 10.1159/000313379.

8. : Royal Botanic Gardens, Kew: Plant DNA C-values database. http://data.kew.org/cvalues/ Accessed 2018 Aug 15.

9. Byrne SL, Nagy I, Pfeifer M, Armstead I, Swain S, Studer B, et al.. A synteny-based draft genome sequence of the forage grass Lolium perenne. Plant J Cell Mol Biol. 2015; doi: 10.1111/tpj.13037.

10. Velmurugan J, Mollison E, Barth S, Marshall D, Milne L, Creevey CJ, et al.. An ultra-high density genetic linkage map of perennial ryegrass (Lolium perenne) using genotyping by sequencing (GBS) based on a reference shotgun genome assembly. Ann Bot. 2016; doi: 10.1093/aob/mcw081.

11. : ASM173568v1 - Genome - Assembly - NCBI. https://www.ncbi.nlm.nih.gov/assembly/GCA_001735685.1/ Accessed 2018 Aug 20.

12. Knorst V, Yates S, Byrne S, Asp T, Widmer F, Studer B, et al.. First assembly of the gene-space of Lolium multiflorum and comparison to other Poaceae genomes. Grassl Sci. doi:10.1111/grs.12225.

13. Huang L, Feng G, Yan H, Zhang Z, Bushman BS, Wang J, et al.. Genome assembly provides insights into the genome evolution and flowering regulation of orchardgrass. Plant Biotechnol J. doi: 10.1111/pbi.13205.

14. Myers EW. Toward Simplifying and Accurately Formulating Fragment Assembly. J Comput Biol. 1995; doi: 10.1089/cmb.1995.2.275.

15. Bredeson JV, Lyons JB, Prochnik SE, Wu GA, Ha CM, Edsinger-Gonzales E, et al.. Sequencing wild and cultivated cassava and related species reveals extensive interspecific hybridization and genetic diversity. Nat Biotechnol. 2016; doi: 10.1038/nbt.3535.

16. Xia E-H, Zhang H-B, Sheng J, Li K, Zhang Q-J, Kim C, et al.. The Tea Tree Genome Provides Insights into Tea Flavor and Independent Evolution of Caffeine Biosynthesis. Mol Plant. 2017; doi: 10.1016/j.molp.2017.04.002.

17. Copetti D, Búrquez A, Bustamante E, Charboneau JLM, Childs KL, Eguiarte LE, et al.. Extensive gene tree discordance and hemiplasy shaped the genomes of North American columnar cacti. Proc Natl Acad Sci. 2017; doi: 10.1073/pnas.1706367114.

18. Edger PP, Poorten TJ, VanBuren R, Hardigan MA, Colle M, McKain MR, et al.. Origin and evolution of the octoploid strawberry genome. Nat Genet. 2019; doi: 10.1038/s41588-019-0356-4.

19. Shirasawa K, Azuma A, Taniguchi F, Yamamoto T, Sato A, Hirakawa H, et al.. De novo whole-genome assembly in interspecific hybrid table grape, ‘Shine Muscat.’bioRxiv. Cold Spring Harbor Laboratory; 2019; doi: 10.1101/730762.

20. Sun X, Jiao C, Schwaninger H, Chao CT, Ma Y, Duan N, et al.. Phased diploid genome assemblies and pan-genomes provide insights into the genetic history of apple domestication. Nat Genet. Nature Publishing Group; 2020; doi: 10.1038/s41588-020-00723-9.

21. Callahan AM, Zhebentyayeva TN, Humann JL, Saski CA, Galimba KD, Georgi LL, et al.. Defining the ‘HoneySweet’ insertion event utilizing NextGen sequencing and a de novo genome assembly of plum (Prunus domestica). Hortic Res. Nature Publishing Group; 2021; doi: 10.1038/s41438-020-00438-2.

22. Kajitani R, Toshimoto K, Noguchi H, Toyoda A, Ogura Y, Okuno M, et al.. Efficient de novo assembly of highly heterozygous genomes from whole-genome shotgun short reads. Genome Res. 2014; doi: 10.1101/gr.170720.113.

23. Safonova Y, Bankevich A, Pevzner PA. dipSPAdes: Assembler for Highly Polymorphic Diploid Genomes. J Comput Biol J Comput Mol Cell Biol. 2015; doi: 10.1089/cmb.2014.0153.

24. Goltsman E, Ho I, Rokhsar D. Meraculous-2D: Haplotype-sensitive Assembly of Highly Heterozygous genomes. ArXiv170309852 Q-Bio. 2017;

25. Weisenfeld NI, Kumar V, Shah P, Church DM, Jaffe DB. Direct determination of diploid genome sequences. Genome Res. 2017; doi: 10.1101/gr.214874.116.

26. Hirsch CN, Hirsch CD, Brohammer AB, Bowman MJ, Soifer I, Barad O, et al.. Draft Assembly of Elite Inbred Line PH207 Provides Insights into Genomic and Transcriptome Diversity in Maize. Plant Cell. 2016; doi: 10.1105/tpc.16.00353.

27. Avni R, Nave M, Barad O, Baruch K, Twardziok SO, Gundlach H, et al.. Wild emmer genome architecture and diversity elucidate wheat evolution and domestication. Science. 2017; doi: 10.1126/science.aan0032.

28. Zhao G, Zou C, Li K, Wang K, Li T, Gao L, et al.. The Aegilops tauschii genome reveals multiple impacts of transposons. Nat Plants. 2017; doi: 10.1038/s41477-017-0067-8.

29. Springer NM, Anderson SN, Andorf CM, Ahern KR, Bai F, Barad O, et al.. The maize W22 genome provides a foundation for functional genomics and transposon biology. Nat Genet. 2018; doi: 10.1038/s41588-018-0158-0.

30. IWGSC TIWGS, Eversole K, Feuillet C, Keller B, Rogers J, Stein N, et al.. Shifting the limits in wheat research and breeding using a fully annotated reference genome. Science. 2018; doi: 10.1126/science.aar7191.

31. Rocha LC, Jankowska M, Fuchs J, Mittelmann A, Techio VH, Houben A. Decondensation of chromosomal 45S rDNA sites in Lolium and Festuca genotypes does not result in karyotype instability. Protoplasma. 2017; doi: 10.1007/s00709-016-0942-6.

32. Luo R, Liu B, Xie Y, Li Z, Huang W, Yuan J, et al.. SOAPdenovo2: an empirically improved memory-efficient short-read de novo assembler. GigaScience. 2012; doi: 10.1186/2047-217X-1-18.

33. Šmarda P, Bureš P, Horová L, Foggi B, Rossi G. Genome Size and GC Content Evolution of Festuca: Ancestral Expansion and Subsequent Reduction. Ann Bot. 2008; doi: 10.1093/aob/mcm307.

34. Singh R, Ming R, Yu Q. Comparative Analysis of GC Content Variations in Plant Genomes. Trop Plant Biol. 2016; doi: 10.1007/s12042-016-9165-4.

35. Mascher M, Gundlach H, Himmelbach A, Beier S, Twardziok SO, Wicker T, et al.. A chromosome conformation capture ordered sequence of the barley genome. Nature. 2017; doi: 10.1038/nature22043.

36. Wu TD, Watanabe CK. GMAP: a genomic mapping and alignment program for mRNA and EST sequences. Bioinforma Oxf Engl. 2005; doi: 10.1093/bioinformatics/bti310.

37. Campbell MS, Law M, Holt C, Stein JC, Moghe GD, Hufnagel DE, et al.. MAKER-P: a tool kit for the rapid creation, management, and quality control of plant genome annotations. Plant Physiol. 2014; doi: 10.1104/pp.113.230144.

38. Pfeifer M, Martis M, Asp T, Mayer KFX, Lübberstedt T, Byrne S, et al.. The perennial ryegrass GenomeZipper: targeted use of genome resources for comparative grass genomics. Plant Physiol. 2013; doi: 10.1104/pp.112.207282.

39. Camacho C, Coulouris G, Avagyan V, Ma N, Papadopoulos J, Bealer K, et al.. BLAST+: architecture and applications. BMC Bioinformatics. 2009; doi: 10.1186/1471-2105-10-421.

40. Quinlan AR, Hall IM. BEDTools: a flexible suite of utilities for comparing genomic features. Bioinformatics. 2010; doi: 10.1093/bioinformatics/btq033.

41. Mollison EMB, Barth S, Milbourne D, Milne L, Halpin C, McCabe M, et al.. De novo Genome Sequencing and Gene Prediction in Lolium perenne, Perennial Ryegrass. Springer Nature;

42. International Brachypodium Initiative. Genome sequencing and analysis of the model grass Brachypodium distachyon. Nature. 2010; doi: 10.1038/nature08747.

43. Jiao Y, Peluso P, Shi J, Liang T, Stitzer MC, Wang B, et al.. Improved maize reference genome with single-molecule technologies. Nature. 2017; doi: 10.1038/nature22971.

44. Simão FA, Waterhouse RM, Ioannidis P, Kriventseva EV, Zdobnov EM. BUSCO: assessing genome assembly and annotation completeness with single-copy orthologs. Bioinformatics. 2015; doi: 10.1093/bioinformatics/btv351.

45. Grabherr MG, Haas BJ, Yassour M, Levin JZ, Thompson DA, Amit I, et al.. Full-length transcriptome assembly from RNA-Seq data without a reference genome. Nat Biotechnol. 2011; doi: 10.1038/nbt.1883.

46. Li H, Durbin R. Fast and accurate long-read alignment with Burrows-Wheeler transform. Bioinforma Oxf Engl. 2010; doi: 10.1093/bioinformatics/btp698.

47. Sim S, Chang T, Curley J, Warnke SE, Barker RE, Jung G. Chromosomal rearrangements differentiating the ryegrass genome from the Triticeae, oat, and rice genomes using common heterologous RFLP probes. TAG Theor Appl Genet Theor Angew Genet. 2005; doi: 10.1007/s00122-004-1916-1.

48. Tang H, Zhang X, Miao C, Zhang J, Ming R, Schnable JC, et al.. ALLMAPS: robust scaffold ordering based on multiple maps. Genome Biol. 2015; doi: 10.1186/s13059-014-0573-1.

49. Novák P, Neumann P, Pech J, Steinhaisl J, Macas J. RepeatExplorer: a Galaxy-based web server for genome-wide characterization of eukaryotic repetitive elements from next-generation sequence reads. Bioinformatics. 29:792–32013;

50. Copetti D, Zhang J, El Baidouri M, Gao D, Wang J, Barghini E, et al.. RiTE database: a resource database for genus-wide rice genomics and evolutionary biology. BMC Genomics. 2015; doi: 10.1186/s12864-015-1762-3.

51. Smit A.F.A., Hubley R., Green P.. RepeatMasker.

52. Krumsiek J, Arnold R, Rattei T. Gepard: a rapid and sensitive tool for creating dotplots on genome scale. Bioinforma Oxf Engl. 2007; doi: 10.1093/bioinformatics/btm039.

53. Jones P, Binns D, Chang H-Y, Fraser M, Li W, McAnulla C, et al.. InterProScan 5: genome-scale protein function classification. Bioinformatics. 2014; doi: 10.1093/bioinformatics/btu031.

54. Kawahara Y, Bastide M de la, Hamilton JP, Kanamori H, McCombie WR, Ouyang S, et al.. Improvement of the Oryza sativa Nipponbare reference genome using next generation sequence and optical map data. Rice. 2013; doi: 10.1186/1939-8433-6-4.

55. Bel MV, Proost S, Wischnitzki E, Movahedi S, Scheerlinck C, Peer YV de, et al.. Dissecting Plant Genomes with the PLAZA Comparative Genomics Platform. Plant Physiol. 2012; doi: 10.1104/pp.111.189514.

56. Peng Z, Lu Y, Li L, Zhao Q, Feng Q, Gao Z, et al.. The draft genome of the fast-growing non-timber forest species moso bamboo (*Phyllostachys heterocycla*). Nat Genet. 2013; doi: 10.1038/ng.2569.

57. Zhang J, Chen L-L, Xing F, Kudrna DA, Yao W, Copetti D, et al.. Extensive sequence divergence between the reference genomes of two elite indica rice varieties Zhenshan 97 and Minghui 63. Proc Natl Acad Sci. 2016; doi: 10.1073/pnas.1611012113.

58. Wang KL-C, Yoshida H, Lurin C, Ecker JR. Regulation of ethylene gas biosynthesis by the *Arabidopsis* ETO1 protein. Nature. 2004; doi: 10.1038/nature02516.

59. Marçais G, Delcher AL, Phillippy AM, Coston R, Salzberg SL, Zimin A. MUMmer4: A fast and versatile genome alignment system. PLOS Comput Biol. 2018; doi: 10.1371/journal.pcbi.1005944.

60. Schnable PS, Ware D, Fulton RS, Stein JC, Wei F, Pasternak S, et al.. The B73 Maize Genome: Complexity, Diversity, and Dynamics. Science. 2009; doi: 10.1126/science.1178534.

61. Chen J, Huang Q, Gao D, Wang J, Lang Y, Liu T, et al.. Whole-genome sequencing of Oryza brachyantha reveals mechanisms underlying Oryza genome evolution. Nat Commun. 2013; doi: 10.1038/ncomms2596.

62. Tang L, Zou X, Achoundong G, Potgieter C, Second G, Zhang D, et al.. Phylogeny and biogeography of the rice tribe (Oryzeae): Evidence from combined analysis of 20 chloroplast fragments. Mol Phylogenet Evol. 2010; doi: 10.1016/j.ympev.2009.08.007.

63. Thomas H, Humphreys MO. Progress and potential of interspecific hybrids of Lolium and Festuca. J Agric Sci. Cambridge University Press; 1991; doi: 10.1017/S0021859600078916.

64. Yamada T, Forster JW, Humphreys MW, Takamizo T. Genetics and molecular breeding in Lolium/Festuca grass species complex. Grassl Sci. 2005; doi: https://doi.org/10.1111/j.1744-697X.2005.00024.x.

65. Kopecký D, Bartoš J, Lukaszewski AJ, Baird JH, Černoch V, Kölliker R, et al.. Development and mapping of DArT markers within the Festuca - Lolium complex. BMC Genomics. 2009; doi: 10.1186/1471-2164-10-473.

66. Humphreys MW. The controlled introgression of Festuca arundinacea genes into Lolium multiflorum. Euphytica. 1989; doi: 10.1007/BF00042621.

67. Kosmala A, Zwierzykowski Z, Gąsior D, Rapacz M, Zwierzykowska E, Humphreys MW. GISH/FISH mapping of genes for freezing tolerance transferred from Festuca pratensis to Lolium multiflorum. Heredity. 2006; doi: 10.1038/sj.hdy.6800787.

