## Supplemental_file_1 for "Evidence for high intergenic sequence variation in heterozygous Italian ryegrass (*Lolium multiflorum* Lam.) genome revealed by a high-quality draft diploid genome assembly"

**Additional file 1**

**Development of a new Italian ryegrass variety, called "Rabiosa".**

The cultivar Rabiosa is derived directly from different ecotype populations collected at various sites in Switzerland (Michaelskreuz, Wiggwil, Gachnang, Root, Beinwil, Bärenbohl) in 1996. Plants from collected seeds were grown in 1997 in isolation fields at Agroscope Reckenholz (Zurich, Switzerland), whereby most persistent plants were allowed to pollinate in 1999 (separate per ecotype population, a schematic representation is given below). In 2000, this improved ecotype offspring was grown in the spaced plant nursery, where in the following year the 31 plants selected for good degree of disease resistance and high vigor were allowed to pollinate (across ecotype populations). Out of these 31 parental plants, offspring of the 24 parents with highest seed yield was grown in another spaced plant nursery trial in 2002. Out of this nursery trial, 11 plants with good performance (disease resistance, vigor) and homogenous flowering time were transplanted into a polycross in 2003. Synthetic-1 (syn-1) seed was harvested from the polycross separately for each of the 11 parental genotypes in 2004. In 2005, 11 half-sib families were then grown in a field experiment with replicated rows of 2.5 m length for the assessment of offspring performance and seed multiplication. Only the 7 best half-sib families were allowed to pollinate (rows of 4 half-sib families eliminated before flowering) and syn-2 seed was harvested in bulk across all rows/half-sib families, forming the base of the cultivar Rabiosa. From this seed batch, one individual seed was germinated, grown and multiplicated for producing plant material for sequencing. This individual was labelled M.02402/16.

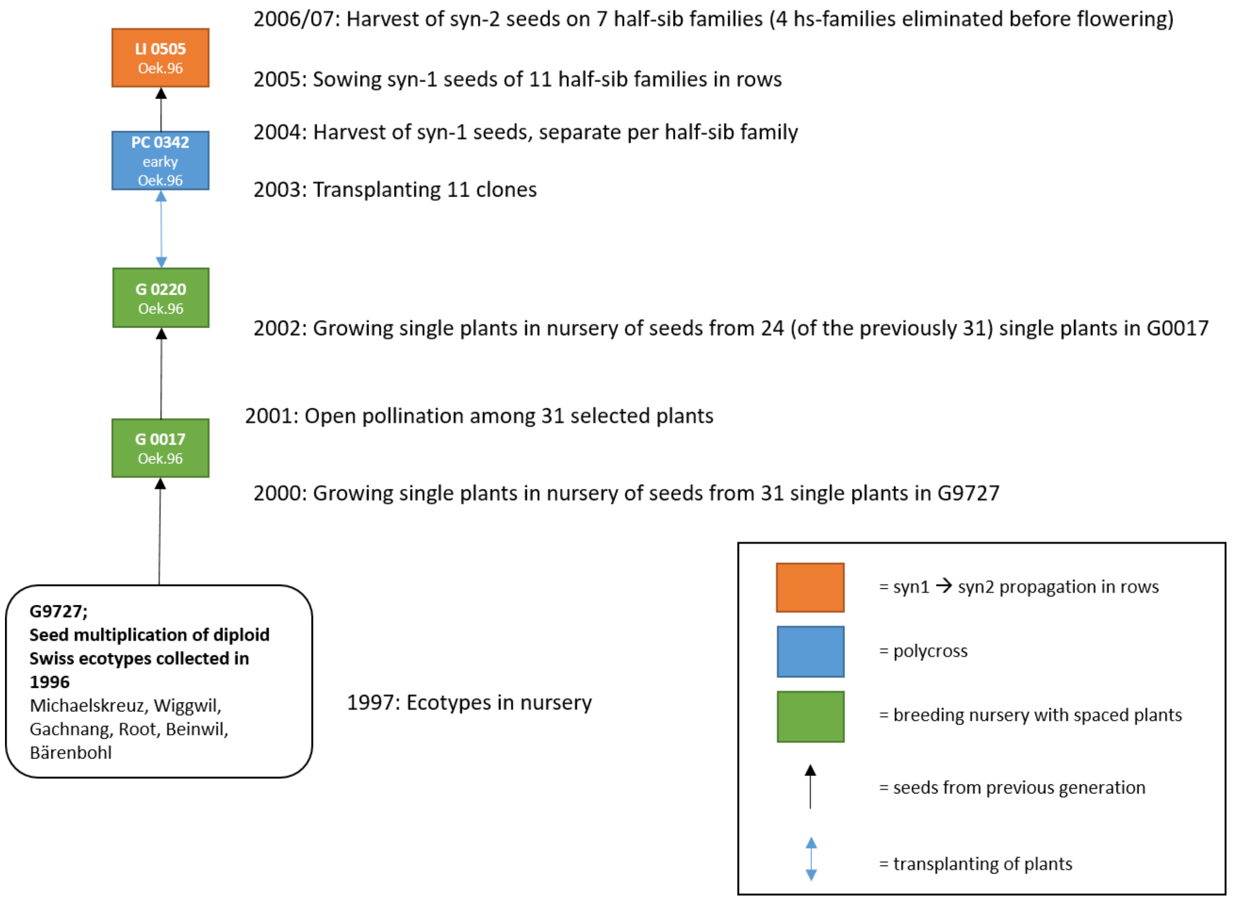
