## Supplemental_file_2 for "Evidence for high intergenic sequence variation in heterozygous Italian ryegrass (*Lolium multiflorum* Lam.) genome revealed by a high-quality draft diploid genome assembly"

**Additional file 2**

### Supplementary methods

#### Cytogenetics

To count chromosome number, cloned tillers of M.02402/16 were transplanted in fresh soil to promote growth of fresh roots. Root tips were sampled, transferred into ice-cold water, and stored in the dark. After 24 h the root tips were soaked in a fixing solution of absolute ethanol and absolute acetic acid (3:1) and stored at 4 °C. Fixed root tips were washed with tap water for 10 min, transferred into 1 N HCl at 60 °C for 10 min, and stained with Schiffs reagent (Carl Roth, Karlsruhe, Germany) for 1 h. Stained root tips were transferred on a glass slide into a drop of acetic acid (30%) and covered with a cover slip. To spread out the cells, covered root tips were squashed by tapping onto the cover slip with a pencil. Chromosomes were visualized on an Olympus BX61 phase contrast microscope, equipped with an ORCA-ER CCD camera (Hamamatsu).

#### Nucleic acid extraction and sequencing

To extract high-quality DNA for sequencing, young leaves were ground in liquid nitrogen to a fine powder, then cells were immediately lysed in a buffer containing 2% CTAB, 0.2 M TrisHCl, 2 M NaCl and 0.05 M EDTA. Lysate was incubated in 0.35 M Sorbitol, 0.1 M Tris HCl and 0.005 EDTA and 0.02 M $Na_2O_5S_2$. DNA was extracted using isopropanol and washed several times using phenol, chloroform and isoamylalcohol (25:24:1). Paired end (PE), PCR-free sequencing libraries were prepared with the TruSeq DNA Sample Preparation Kit version 2 according to the manufacturer's protocol (Illumina, San Diego, CA). Mate pair (MP) libraries were prepared using the Illumina Nextera Mate-Pair Sample Preparation Kit (Illumina, San Diego, CA). The 10x Chromium libraries were prepared following a standard protocol on the Chromium instrument (10x Genomics, Pleasanton, CA). WGS and 10x Chromium libraries were sequenced at different read lengths with Illumina technology (Suppl. Table 1). PE and MP libraries construction and sequencing were conducted at Roy J. Carver Biotechnology Center, University of Illinois at Urbana-Champaign, while 10x Chromium library construction and sequencing were conducted at HudsonAlpha Institute for Biotechnology, Huntsville, Alabama.

The mRNA sequence data used for genome annotation was derived from clones of the same genotype. Four tissue types were sampled: mature leaves, apical meristems, roots and flowers. For mature leaf, two to three 3 cm-long segments were sampled from independent leaves where the ligules had formed. Leaf meristems were harvested by carefully removing all developed leaves then extracting the lowest 3 cm of two growing leaves. Roots were harvested from those protruding outside of the growing pots (approximately 50 mg per sample). Three to four flowers, prior to anthesis, were removed for flower samples. For each organ, three replicate samples were harvested from independent clonally propagated plants and immediately frozen in liquid nitrogen. Total RNA was extracted using TRIzol™ (Thermo Fisher Scientific, Waltham, USA) as described by the manufacturer. The RNA was quality checked using TapeStation 2200 (Agilent, Santa Clara, USA). Sequencing libraries were prepared using Illumina TruSeq preparation, the samples were then sequenced at both ends for 150 bp with Illumina HiSeq2500 and NextSeq at Functional Genomics Center Zurich, Zurich, Switzerland.

### Assembly quality control

Four sets of evidence were used: i) *Lolium* expressed sequence tags (ESTs) and Unigenes were downloaded from NCBI (December 6, 2017) and assembled with Cap3 (v. 02/10/15, `-p 95` [1]) to reduce redundancy. The resulting non-transposable element (TE) transcripts longer than 100 bp were aligned to the scaffolds with GMAP (v. 2017-09-30 [2]) using default parameters. The same unigenes were then aligned to the barley proteome with BLASTP (v. v. 2.5.0 [3]). Only hits longer than 100 aa, covering more than 80% of the query length and with similarity higher than 80% were retained. Hits were manually inspected to remove secondary or aspecific alignments, and remaining alignments were used to link M.02402/16 scaffolds to barley genes (assigned to the 7 chromosome pseudomolecules Additional File 2, sheet Unigenes). ii) Bowtie and Cufflinks [4,5] were run with *Lolium perenne* RNA-Seq data (Studer *et al.*, unpublished) to produce provisional gene models that were later polished by running them through the MAKER (v. 2.31.9 [6]) pipeline. After removing models with similarity to TE genes, the remaining sequences were aligned to the barley proteome as above. After parsing spurious or multiple hits, 7082 gene alignments were connecting 1435 M.02402/16 scaffolds to barley gene models anchored on chromosomes (Additional File 2, sheet RNA-Seq). iii) A gene-guided whole-genome alignment was performed in CoGe (https://genomevolution.org/coge/). Collinear blocks of three or more genes were computed with SynMap (Additional File 2, sheet SynMap). iv) the 762 markers of the *L. perenne* genetic map [7] were aligned to the assembly with BLASTN, and hits covering at least 90% of query length with at least 90% similarity were retained. Using this genetic evidence from a closely-related species, the genetic markers aligned 411 scaffolds to *L. perenne* linkage groups (LGs, Additional File 2, sheet Pfeifer 2013).

Candidate chimeric scaffolds were further inspected with the following steps. Bowtie2 (v. 2.2.3, with the `--very-sensitive` option and appropriate insert size for each library [4]) was used to align the 6 and 9 kb MP data. The alignments were parsed with SAMtools view and sort functions (v. 0.1.19 [8]) to retain only pairs that aligned to the same scaffold. Tigmint (v. 1.1.2 [9]) was run to map the 10x linked reads and infer the extents of the DNA molecules on the assembly.

### Completeness

To assess the completeness of the assembly, the BUSCO (v. 3 [10]) analysis was run in genome mode with wheat as a species for Augustus with the Viridiplantae dataset.
To estimate the fraction of gene space that is missing from the assembly, the raw RNA-Seq reads were aligned to the M.02402/16 reference with STAR (v. 2.5.3a, `--outReadsUnmapped Fastx --alignIntronMax 10000 --outSAMstrandField intronMotif --outSAMattrIHstart 0 --outSAMtype BAM SortedByCoordinate` [11]). Upon assembly with Trinity (v. 2.5.1 [12]), *de novo* transcripts were aligned to the reference with GMAP (v. 2018-03-25, `--max-intronlength-middle=10000 -L 50000 --no-chimeras --min-trimmed-coverage=0.7 --min-identity=0.90 -f psl -n 2 -Y`).

After removing low quality bases and short reads with Trimmomatic (v. 0.36 `ILLUMINACLIP:TruSeq3-PE.fa:2:30:10 SLIDINGWINDOW:4:30 MINLEN:50` [13], PE470 reads were clipped to 200 bp, `CROP:200`), PE reads were aligned to the assembly with BWA-MEM (v. 0.7.17, optimizing the size parameters for each library [14]). SAMtools fixmate, sort, merge, depth (v. 0.1.19 [8]) were used to remove secondary alignments and unaligned reads.

The K-mer Analysis Toolkit (KAT v.2.3.1, `kat comp -m 23` [15]) was run with the PE700 library and the final assembly.

#### Repeat annotation

The RepeatExplorer (v. v0.2.8-2479 [16]) and RepeatModeler [17] pipelines were run at default settings. After reducing the redundancy with vsearch (v 2.7.0, `--id 0.95 --strand both –centroids` [18]), the resulting centroid sequences were characterized according to [19] by comparing them to the TREP database (v. 17, August 2016 [20]). Sequences with significant hits to non-TE proteins in the NCBI nr database were removed. Repeats and TEs in the assembly were detected with RepeatMasker (v. open-4.0.6 [17]) adopting this custom library. TE coding sequences were identified by homology using BLASTER and MATCHER (v2.5 [21]) and later the nucleotidic and protein TE annotations were reconciled as in [22].

#### Protein-coding genes and non-coding RNA annotation

Gene models were derived in scaffolds longer than 5 kb by running them through MAKER pipeline. Sets of evidence were derived from *Lolium* EST-derived unigenes (see ***Assembly quality control and sequence anchoring*** in the main body), clustered (VSEARCH v 2.7.0 `--id 0.90 –centroids` [18]) barley [23], rice [24], and Arabidopsis [25] proteomes, and gene models from RNA-Seq data. *L. multiflorum* (SRA accession number SRP151232), *L. perenne* (Studer et al., unpublished) and other transcripts ([26], Suppl. Table 2) were first trimmed with fastp (v 0.12.4, `–l 36 –c –g –p –M 30 –5 –3` [27]), then aligned with STAR. Alignment files were merged by species or sample (SAMtools merge), and transcripts were predicted with StringTie (v 1.3.3b, `–m 100` [28]). The *ab initio* gene prediction was performed with custom-trained Augustus (v. 3.2.3 [29]) and SNAP (v. 2006-07-28 [30]) hmm files according to [31]. Predicted protein-coding loci were removed if they contained Gene Ontology domains with homology to TE coding regions (IPR000477, IPR001207, IPR001584, IPR002559, IPR004242, IPR004252, IPR004264, IPR004330, IPR004332, IPR005063, IPR005162, IPR006912, IPR007321, IPR013103, IPR013242, IPR014736, IPR015401, IPR018289, IPR026103, IPR026960, IPR027806, upon analysis with InterProScan v.5.31-70.0 [32]). Also, models were removed if at least 40% of their sequence (or at least 100 bp) aligned at more than 40% similarity to a TE coding region (BLASTP), or if CDS exons were overlapping >80% of their length with interspersed TE repeats from RepeatMasker.
The functional annotation was performed via BLASTP (`–evalue 1e-15`) alignment to Arabidopsis proteins and with InterProScan 5 (using default setting and databases). For the former method, the protein sequences were aligned (against Arabidopsis proteins [25] and the best hit was used as the annotation source. A Perl script was then used to retrieve the gene name and Gene Ontology (GO) terms from the corresponding Arabidopsis proteins. For each gene model, the resulting InterProScan matches were then collapsed and combined with the Arabidopsis data.
CoreGF (v 2.5 [33]) was run with the PLAZA2.5 monocot dataset, and BUSCO (v3 [10], Suppl. Table 5) was un in proteome mode with the same datasets as above.
Non-coding RNAs were annotated with tRNA-Scan SE (v 2.0, [34]) and Infernal (v 1.1.2, `--cut_ga --rfam --nohmmonly --fmt 2 --oskip --oclan` [35]), adopting the Rfam database (v. 13.0 [36]). Infernal's output was parsed to remove duplicated predictions.

#### Sequence diversity in allelic regions

Gene alignments were performed with GMAP (v. 2018-03-25, `--max-intronlength-middle=10000 -L 50000 --no-chimeras --min-trimmed-coverage=0.7 --min-identity=0.90 --cross-species -n 2 -Y`). To collinear blocks were identified via BLASTP of barley and Rabiosa proteomes and the results were processed by MCSCanX [37] and visualized with VGSC (v1.1 [38]) the dotplot alignments were generated with Gepard (v. 1.40 [39]). Scaffolds were aligned with MUMmer4.x utilities (`nucmer -L 500 -c 100`, `delta-filter -1`, `dnadiff` [40]).

### References


1. Huang X, Madan A. CAP3: A DNA Sequence Assembly Program. *Genome Res*. 1999; doi: 10.1101/gr.9.9.868.

2. Wu TD, Watanabe CK. GMAP: a genomic mapping and alignment program for mRNA and EST sequences. *Bioinforma Oxf Engl*. 2005; doi: 10.1093/bioinformatics/bti310.

3. Camacho C, Coulouris G, Avagyan V, Ma N, Papadopoulos J, Bealer K, et al.. BLAST+: architecture and applications. *BMC Bioinformatics*. 2009; doi: 10.1186/1471-2105-10-421.

4. Langmead B, Salzberg SL. Fast gapped-read alignment with Bowtie 2. *Nat Methods*. 2012; doi: 10.1038/nmeth.1923.

5. Trapnell C, Roberts A, Goff L, Pertea G, Kim D, Kelley DR, et al.. Differential gene and transcript expression analysis of RNA-seq experiments with TopHat and Cufflinks. *Nat Protoc*. 2012; doi: 10.1038/nprot.2012.016.

6. Campbell MS, Law M, Holt C, Stein JC, Moghe GD, Hufnagel DE, et al.. MAKER-P: a tool kit for the rapid creation, management, and quality control of plant genome annotations. *Plant Physiol*. 2014; doi: 10.1104/pp.113.230144.

7. Pfeifer M, Martis M, Asp T, Mayer KFX, Lübberstedt T, Byrne S, et al.. The perennial ryegrass GenomeZipper: targeted use of genome resources for comparative grass genomics. *Plant Physiol*. 2013; doi: 10.1104/pp.112.207282.

8. Li H. A statistical framework for SNP calling, mutation discovery, association mapping and population genetical parameter estimation from sequencing data. *Bioinforma Oxf Engl*. 2011; doi: 10.1093/bioinformatics/btr509.

9. Jackman SD, Coombe L, Chu J, Warren RL, Vandervalk BP, Yeo S, et al.. Tigmint: Correcting Assembly Errors Using Linked Reads From Large Molecules. *bioRxiv*. 2018; doi: 10.1101/304253.

10. Simão FA, Waterhouse RM, Ioannidis P, Kriventseva EV, Zdobnov EM. BUSCO: assessing genome assembly and annotation completeness with single-copy orthologs. *Bioinformatics*. 2015; doi: 10.1093/bioinformatics/btv351.

11. Dobin A, Davis CA, Schlesinger F, Drenkow J, Zaleski C, Jha S, et al.. STAR: ultrafast universal RNA-seq aligner. *Bioinforma Oxf Engl*. 2013; doi: 10.1093/bioinformatics/bts635.

12. Grabherr MG, Haas BJ, Yassour M, Levin JZ, Thompson DA, Amit I, et al.. Full-length transcriptome assembly from RNA-Seq data without a reference genome. *Nat Biotechnol*. 2011; doi: 10.1038/nbt.1883.

13. Bolger AM, Lohse M, Usadel B. Trimmomatic: a flexible trimmer for Illumina sequence data. *Bioinforma Oxf Engl*. 2014; doi: 10.1093/bioinformatics/btu170.

14. Li H, Durbin R. Fast and accurate long-read alignment with Burrows-Wheeler transform. *Bioinforma Oxf Engl*. 2010; doi: 10.1093/bioinformatics/btp698.

15. Mapleson D, Garcia Accinelli G, Kettleborough G, Wright J, Clavijo BJ, Berger B. KAT: a K-mer analysis toolkit to quality control NGS datasets and genome assemblies. *Bioinformatics*. 2017; doi: 10.1093/bioinformatics/btw663.

16. Novák P, Neumann P, Pech J, Steinhaisl J, Macas J. RepeatExplorer: a Galaxy-based web server for genome-wide characterization of eukaryotic repetitive elements from next-generation sequence reads. *Bioinformatics*. 29:792–7932013;

17. Smit A.F.A., Hubley R., Green P.. RepeatMasker.

18. Rognes T, Flouri T, Nichols B, Quince C, Mahé F. VSEARCH: a versatile open source tool for metagenomics. *PeerJ*. 2016; doi: 10.7717/peerj.2584.

19. Wicker T, Sabot F, Hua-Van A, Bennetzen JL, Capy P, Chalhoub B, et al.. A unified classification system for eukaryotic transposable elements. *Nat Rev Genet*. 2007; doi: 10.1038/nrg2165.

20. Wicker T, Matthews DE, Keller B. TREP: a database for Triticeae repetitive elements. *Trends Plant Sci*. 2002; doi: 10.1016/S1360-1385(02)02372-5.

21. Flutre T, Duprat E, Feuillet C, Quesneville H. Considering Transposable Element Diversification in De Novo Annotation Approaches. *PLoS ONE*. 2011; doi: 10.1371/journal.pone.0016526.

22. Stein JC, Yu Y, Copetti D, Zwickl DJ, Zhang L, Zhang C, et al.. Genomes of 13 domesticated and wild rice relatives highlight genetic conservation, turnover and innovation across the genus Oryza. *Nat Genet*. 2018; doi: 10.1038/s41588-018-0040-0.

23. Mascher M, Gundlach H, Himmelbach A, Beier S, Twardziok SO, Wicker T, et al.. A chromosome conformation capture ordered sequence of the barley genome. *Nature*. 2017; doi: 10.1038/nature22043.

24. Kawahara Y, Bastide M de la, Hamilton JP, Kanamori H, McCombie WR, Ouyang S, et al.. Improvement of the Oryza sativa Nipponbare reference genome using next generation sequence and optical map data. *Rice*. 2013; doi: 10.1186/1939-8433-6-4.

25. Lamesch P, Berardini TZ, Li D, Swarbreck D, Wilks C, Sasidharan R, et al.. The Arabidopsis Information Resource (TAIR): improved gene annotation and new tools. *Nucleic Acids Res*. 2012; doi: 10.1093/nar/gkr1090.

26. Pan L, Zhang X, Wang J, Ma X, Zhou M, Huang L, et al.. Transcriptional Profiles of Drought-Related Genes in Modulating Metabolic Processes and Antioxidant Defenses in Lolium multiflorum. *Front Plant Sci*. 2016; doi: 10.3389/fpls.2016.00519.

27. Chen S, Zhou Y, Chen Y, Gu J. fastp: an ultra-fast all-in-one FASTQ preprocessor. *bioRxiv*. 2018; doi: 10.1101/274100.

28. Pertea M, Kim D, Pertea GM, Leek JT, Salzberg SL. Transcript-level expression analysis of RNA-seq experiments with HISAT, StringTie and Ballgown. *Nat Protoc*. 2016; doi: 10.1038/nprot.2016.095.

29. Stanke M, Waack S. Gene prediction with a hidden Markov model and a new intron submodel. *Bioinformatics*. 2003; doi: 10.1093/bioinformatics/btg1080.

30. Korf I. Gene finding in novel genomes. *BMC Bioinformatics*. 2004; doi: 10.1186/1471-2105-5-59.

31. Bowman MJ, Pulman JA, Liu TL, Childs KL. A modified GC-specific MAKER gene annotation method reveals improved and novel gene predictions of high and low GC content in Oryza sativa. *BMC Bioinformatics*. 2017; doi: 10.1186/s12859-017-1942-z.

32. Jones P, Binns D, Chang H-Y, Fraser M, Li W, McAnulla C, et al.. InterProScan 5: genome-scale protein function classification. *Bioinformatics*. 2014; doi: 10.1093/bioinformatics/btu031.

33. Bel MV, Proost S, Wischnitzki E, Movahedi S, Scheerlinck C, Peer YV de, et al.. Dissecting Plant Genomes with the PLAZA Comparative Genomics Platform. *Plant Physiol*. 2012; doi: 10.1104/pp.111.189514.

34. Schattner P, Brooks AN, Lowe TM. The tRNAscan-SE, snoscan and snoGPS web servers for the detection of tRNAs and snoRNAs. *Nucleic Acids Res*. 2005; doi: 10.1093/nar/gki366.

35. Nawrocki EP, Eddy SR. Infernal 1.1: 100-fold faster RNA homology searches. *Bioinforma Oxf Engl*. 2013; doi: 10.1093/bioinformatics/btt509.

36. Kalvari I, Argasinska J, Quinones-Olvera N, Nawrocki EP, Rivas E, Eddy SR, et al.. Rfam 13.0: shifting to a genome-centric resource for non-coding RNA families. *Nucleic Acids Res*. 2018; doi: 10.1093/nar/gkx1038.

37. Wang Y, Tang H, DeBarry JD, Tan X, Li J, Wang X, et al.. MCScanX: a toolkit for detection and evolutionary analysis of gene synteny and collinearity. *Nucleic Acids Res*. 2012; doi: 10.1093/nar/gkr1293.

38. Xu Y, Bi C, Wu G, Wei S, Dai X, Yin T, et al.: VGSC: A Web-Based Vector Graph Toolkit of Genome Synteny and Collinearity. BioMed Res. Int. https://www.hindawi.com/journals/bmri/2016/7823429/ (2016). Accessed 2019 Aug 9.

39. Krumsiek J, Arnold R, Rattei T. Gepard: a rapid and sensitive tool for creating dotplots on genome scale. *Bioinforma Oxf Engl*. 2007; doi: 10.1093/bioinformatics/btm039.

40. Marçais G, Delcher AL, Phillippy AM, Coston R, Salzberg SL, Zimin A. MUMmer4: A fast and versatile genome alignment system. *PLOS Comput Biol*. 2018; doi: 10.1371/journal.pcbi.1005944.
